# Isolation of stage-specific spermatogenic cells by dynamic histone incorporation and removal in spermatogenesis

**DOI:** 10.1101/2023.06.25.546481

**Authors:** Yasuhiro Fujiwara, Masashi Hada, Yuko Fukuda, Chizuko Koga, Erina Inoue, Yuki Okada

**Affiliations:** Institute of Quantitative Biosciences, The University of Tokyo, 1-1-1 Yayoi, Bunkyo-ku, Tokyo 113-0032, Japan

**Keywords:** spermatogenesis, fluorescence-activated cell sorting, histones, fluorescence tagging, testicular germ cells

## Abstract

Due to the lack of an *in vitro* spermatogenesis system, studies on mammalian spermatogenesis require the isolation of specific germ cell populations for further analyses. BSA gradient and elutriation have been used for several decades to purify testicular germ cells; more recently, fluorescence-activated cell sorting (FACS) has become popular. Although each method has its advantages and disadvantages and is used depending on the purpose of the experiment, reliance on FACS is expected to be more prevalent because fewer cells can be managed. However, the currently used FACS method for testicular germ cells relies on karyotypic differences via DNA staining. Thus, it remains challenging to separate post-meiotic haploid cells (spermatids) according to their differentiation stage despite significant variations in morphology and chromatin state. In this study, we developed a method for finely separating testicular germ cells using VC mice carrying fluorescently tagged histones. This method enables the separation of spermatogonia, spermatocytes, and spermatids based on the intensity of histone fluorescence and cell size. Combined with a DNA staining dye, this method separates spermatids after elongation according to each spermiogenic stage. Although the necessity for a specific transgenic mouse line is less versatile, this method is expected to be helpful for the isolation of testicular germ cell populations because it is highly reproducible and independent of complex cell sorter settings and staining conditions.

## Introduction

Sperm cells are produced through a complex developmental process called spermatogenesis; the most notable dynamic change during this process is in the DNA content associated with epigenetic modifications (1–3). In mammals, postnatal spermatogenesis begins with unipotent tissue-specific stem cells, designated as spermatogonial stem cells (SSCs), containing a 2N nucleus. SSCs then differentiate into spermatogonia through repeated somatic-type cell divisions. These differentiated spermatogonia then replicate their DNA content to 4N to initiate meiosis I. Phase I (prophase I) takes approximately 10 days in mice and is divided into substages according to the morphology of the axial elements: leptotene, zygotene, pachytene, and diplotene. After two rounds of meiotic chromosomal segregations via a transient cell status called secondary spermatocytes, the entire process of meiosis can be completed within 24 h. The resulting haploid spermatids (1N) undergo further differentiation, called spermiogenesis, which consists of 16 steps in mice (4,5). At approximately step 9, spermatids start to pack their chromatin by replacing the majority of their histones with transition proteins (TNPs) and protamines (PRMs), concomitant with massive histone degradation (6,7).

Various approaches have been used to isolate testicular germ cells at specific spermatogenic stages, such as centrifugal elutriation (8,9), BSA gradient sedimentation/STA-PUT (10–14), and, more recently, fluorescence-activated cell sorting (FACS) (15–19). One of the advantages of centrifugal elutriation and BSA gradient sedimentation is that they can manage a large number of cells (>10^8^ cells) within a few hours, with the purity of the isolated cells usually maintained at approximately 70-90%. In contrast, FACS can achieve higher purity (>95%), although it cannot handle many cells (<10^8^ cells) in a relatively shorter period of time. The recent trend toward more sensitive experimental analyses and an increase in the number of purity-oriented experiments, such as next-generation sequencing, will likely lead to more opportunities to use FACS for isolating testicular germ cells.

Currently, conventional FACS methods for testicular germ cells are exclusively DNA content-dependent and use DNA-staining dyes such as Hoechst 33342 and Dye Cycle Violet (DCV) have been demonstrated to be particularly useful for the separation of cells across meiotic subpopulations, such as leptotene, zygotene, pachytene, and diplotene cells (15,16,19,20). In contrast, post-meiotic cells (e.g., spermatids) are recognized by these stains as a single population. More recently, SYTO16 can be applied to the fine separation of spermatids because of its unique high affinity for spermatids at the later stages of differentiation (21,22). In fixed cells, SYTO16 staining can separate spermatids into four populations (steps 1–9, 10–12, 13–14, and 15–16), each with an extremely high purity of 95–100% (21). In living cells, on the other hand, SYTO16 can separate three populations (round, early elongating, and late elongating spermatids); the latter two populations have been shown to be 10–20% cross-contaminated with each other (22).

We previously generated a transgenic mouse line (named “VC mouse”) that carries two transgenes, H4-Venus (H4V) and H3.3-mCherry (H3.3C), in their testicular germ cells (23). Histologically, these transgenes are exclusively expressed in a differentiation stage-specific manner; H4V is expressed in spermatogonia, spermatocytes, and round spermatids with different intensities in Venus, whereas H3.3C is expressed in elongating and elongated spermatids (23). To take advantage of this property, we established a unique method to divide spermatogenic germ cell populations using FACS according to the expression levels of H4V and H3.3C in combination with cell size. This method allows testicular germ cell populations with the same DNA content, especially post-meiotic spermatids, to be separated into subpopulations without the need for immunostaining in a stepwise manner.

## Materials and methods

### Mice

The homozygous H4-Venus/H3.3-mCherry (VC) transgenic mice were maintained in C57BL6/J background in the mouse facility of the institute. Three-to eight-month-old male mice were used for experiments except for isolation of meiotic cells, in which 4-5-week-old male mice were used. Animal experiments in this study were approved by the Institutional Animal Care and Use Committee (approval #0212, 0313, 0409). All methods and animal protocols were carried out in accordance with institutional animal guidelines and regulations.

### Preparation of single cell suspension from mouse testis

The tunica albuginea was removed from testes. The testes were then gently loosened with tweezers and transferred to PBS containing 0.5 mg/ml Collagenase type I (Sigma Aldrich). Tissues were incubated at 37□C for 5-10 min until the seminiferous tubules were dispersed. After brief centrifugation, the supernatant was discarded and 0.5 % Trypsin (Thermo-Fisher) containing 2 mg/ml DNase I (Sigma Aldrich) was added to the pellet. The pellet was briefly pipetted up and down and incubate at 37□C for digestion. Pipetting was performed every 5 min until no obvious cell clumps was visible (usually up to 15 min). One third volume of FBS was then added to inactivate the trypsin. Cells were filtered through a 70 μm cell strainer and washed with PBS containing 2% FBS three times. For DNA staining, Vybrant DyeCycle Violet Stain (Themo Fisher) was added at the final concentration of 1 μM followed by incubation at 37□C for 30 min in dark. Cells were filtered again through a 50 μm cell strainer immediately before sorting.

### Antibody Staining for flowcytometry analyses

Anti-CD9-Alexa 647 (clone MZ3) (BioLegend, #124809), anti-cKit-APC (clone ACK2) (TONBO Bioscience, #20-1172), anti-MCAM-APC (clone MF-9F1) (BioLegend, #134712), rat Isotype IgG2a-APC (clone LTF2) (TONBO Bioscience, #20-4031) were added to single cell suspension with 1:100 dilution. Staining was performed for 45-60 min on ice.

### Cell Sorting

A BD FACS Aria III was used for cell sorting. Data acquisition was performed using BD FACSDiva software. 100 μm nozzle was used. 405, 488, 561, and 633 nm lasers were used. Sort mode was set to 4 Way Purity with flow rate as 1.0. The filter sets used for sorting and detection were as follows: i) DCV: 450/40, ii) Venus: 502LP/530/30, iii) mCherry: 600LP/610/20, iv) APC: 660/20. Settings of working plots were indicated in Results. Flowjo (FlowJo LLC) and NovoExpress (Agilent Technologies) softwares were used for data analyses and depiction.

### Immunostaining, image acquisition, and analyses

Sorted cells were applied onto MAS-or poly-A-lysine-coated multi-spot slide glasses (Matsunami glass). Cells were left for 15-30 min to allow them attached to the slide glass, then fixed with 1 % paraformaldehyde. After PBS wash and blocking, following primary antibodies were applied; anti-γH2AX antibody (abcam #22551, 1:2,000), rat anti-SYCP3 antibody (1:1,000) (24) follower by applying the Alexa fluorescence-labeled secondary antibodies (Thermo Fisher). PNA-Cy5 (Vector laboratories) and Hoechst 33342 (Thermo Fisher) were used for counterstaining and histological staging of seminiferous tubules. Olympus IX83 microscope (Evident Japan), a digital CMOS camera ORCA-spark (Hamamatsu Photonics), cellSens software (Evident Japan) were used for image acquisition, and the obtained images were processed using ImageJ software. All the images used in the figures were the representative result, and each experiment were repeated at least three times.

### Elongated spermatid injection (ELSI)

For oocyte collection, C57BL6/J female mice were injected intraperitoneally with 5 units of pregnant mare serum gonadotropin (ASKA Pharmaceutical) and 50 μL of anti-inhibin serum (Central research), followed 48-50 hours later by an injection of 7.5 units of human chorionic gonadotropin (hCG) (ASKA Pharmaceutical) (25). MII oocytes were collected from the oviducts 15-17 hours after hCG treatment, and cumulus cells were removed by treatment for 5 minutes with 48 units/mL hyaluronidase (FUJIFILM Irvine Scientific) diluted in HTF medium (ARK Resource). The oocytes were transferred to a fresh HTF medium and incubated for 1 hour at 37°C in an atmosphere of 5% CO2. Testis single-cell suspensions from VC mice were subjected to FACS without DCV staining. Condensed spermatids were sorted and used for microinjection. The cell membrane of typical condensed spermatids was broken using a pneumatic microinjector (Narishige), and the nuclei were injected into oocytes using a micropipette attached to a Piezo-electric actuator (Prime Tech). Surviving oocytes were cultured in HTF medium at 37□C overnight. Embryos that reached the 2-cell stage were transferred into the uteri of pseudopregnant ICR female mice.

## Results

### Tracking of spermatogenic cell differentiation using VC mouse testes

Representative results of the sorting of testicular cells isolated from adult VC male mice were shown in Fig. 1. Presumably, non-living cells in the range of [FSC<20×10^3^/SSC>15×10^3^] were first eliminated through gating, while cells within the range [FSC<20×10^3^/SSC≤15×10^3^] were intentionally retained considering the small size of their condensed spermatids (Fig. 1A). After gating singlet cells (Fig. 1B), DCV positive (DCV+) cells were gated (Fig. 1C). When the DCV+ population was plotted according to FSC/SSC, three major cell populations, designated small, middle, and large, were identified (Fig. 1D). DNA content analyses revealed that the small cells mainly constituted the left-shifted 1C, medium-sized cells constituted the 1C, and large cells constituted the 2C and 4C peaks (Fig. 1E, 1F).

**Figure 1.**
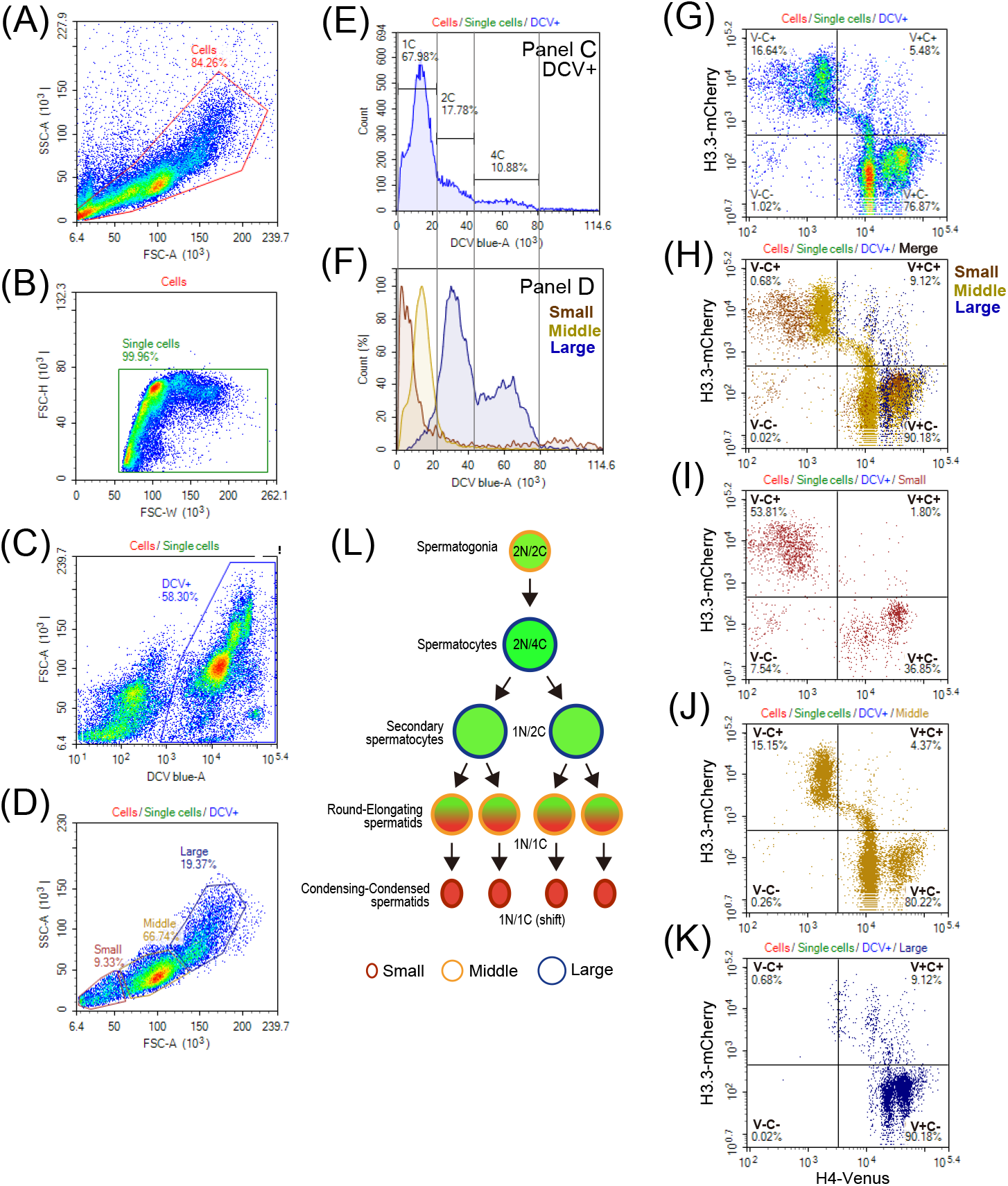
Gating strategy to identify testicular germ cells from VC mice. (A) Removal of non-cellular contaminants. (B) Removal of non-single cells. (C) Gating of DCV+ cells. (D) Setting of gates according to cell size. (E) DNA content analysis of DCV+ cells. (F) DNA content analysis of small, middle, and large-sized cells. (G) Expression pattern of Venus (H4V, X-axis) and mCherry (H3.3C, Y-axis) in DCV+ cells. (H–K) Expression pattern of H4V and H3.3C in all merged (H), small (I), middle (J), and large (K)-sized cells. (L) Schematic summary of the sorting strategy for testicular cells in VC mice depending on cell size, nuclear content, and the expression of H4V and H3.3C.

The DCV+ population was further examined in H4-Venus and H3.3-mCherry plots. Consistent with a previous report (23), the cells were continuously aligned from the Venus+/mCherry-(V+C-) to the Venus-/mCherry+ (V-C+) population (Fig. 1G). The plot in Fig. 1H also shows the characteristic distribution of the cells of each size. First, >50 % of the small cells were plotted in the V-C+ area (Fig. 1I). As their DNA content exhibited a left-shift 1C (Fig. 1F, brown line), they were predicted to be condensing-condensed spermatids. Second, the middle-sized cells included a variety of cell types with dynamically variable Venus and mCherry expression levels (Fig. 1J), whose DNA content was predominantly 1C, with a few 2C (Fig. 1F, yellow line). Because fluorescent microscopic analyses revealed that a color change from Venus to mCherry occurred in step 8–11 spermatids (23), an intermediate, continuous population in the V+C+ areas likely represented step 9–10 spermatids. The other middle-sized cell population had higher H4V expression in the V+C-area, which likely consisted of spermatogonia because of the similarity of their cell size to round spermatids (Fig. 1J). Finally, the large cells were mainly plotted in the V+C-area and formed two adjacent populations with different H4V expression levels (Fig. 1K); the DNA content of these cells was 2C and 4C (Fig. 1F, blue line). Based on these observations, combined with the reported histological features of VC testes (23), the predicted cell properties are summarized in Fig. 1L.

Since DCV-negative (DCV-) substances were observed at non-negligible levels (Fig. 1C, S1A), their properties were also examined. We found that they consisted mainly of smaller-sized cells and were highly concentrated in the V-C+ areas in the Venus/mCherry plots; some of them continuously expanded into the V-C-areas (Fig. S1B–C). Based on these observations, it was inferred that these substances were residual bodies derived from condensing and condensed spermatids containing H4V and H3.3C remnants that underwent degradation.

### Identification of spermatogonial cell populations

Next, we attempted to confirm whether the middle-sized cells with high Venus expression were spermatogonia (Fig. 2A, a black square). To identify their properties, the cells were co-stained for MCAM (broad spermatogonial marker) (26), CD9 (undifferentiated spermatogonial marker) (27), and c-Kit (differentiated spermatogonial marker) (28). We found that ∼98% of the cells were MCAM-positive (Fig. 2B). In addition, ∼84% and 45% of them expressed CD9 and c-Kit, respectively (Fig. 2C, 2D), supporting our prediction that this population consisted of spermatogonia.

**Figure 2.**
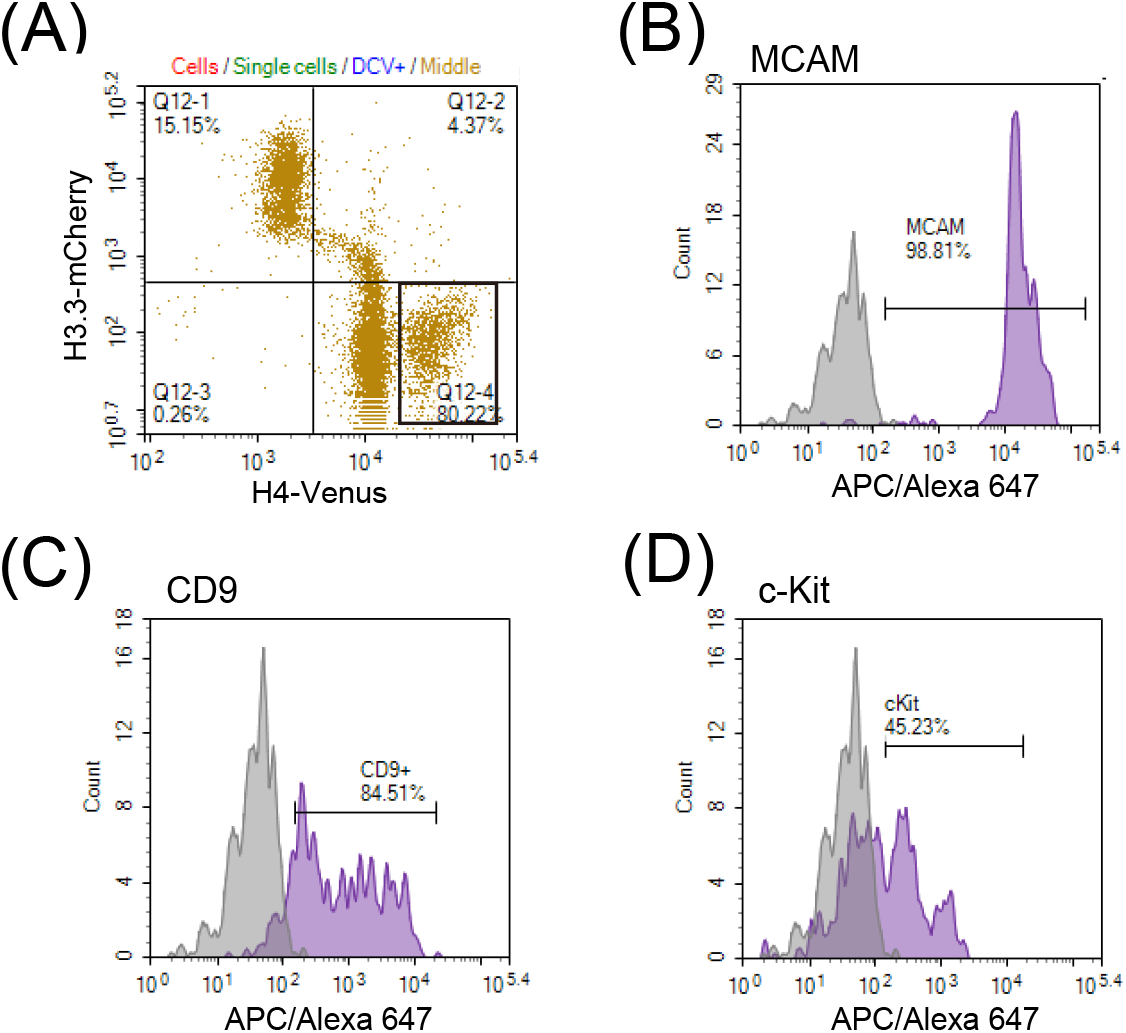
Characterization of spermatogonia. (A) Expression of H4V and H3.3C in middle-sized cells (this plot is the same as Fig. 1J). A cell population suspected to be spermatogonia is surrounded by a black square. (B–D) Expression of MCAM (B), CD9 (C), and c-Kit (D) in the spermatogonial cell population. Gray histograms indicate isotype controls.

### Isolating subpopulations of spermatocytes

Although isolating spermatocyte subpopulations via FACS using DNA staining dyes is well-established (15,16,19,20), we also tested the utility of FACS in separating spermatocyte subpopulations in VC testes. To reduce contamination with post-meiotic cells, 4-5-week-old testes were used. MCAM and DCV staining were then performed to distinguish spermatogonia and examine the DNA content, respectively. Middle- and large-sized cells were gated and DCV+ cells were stained with MCAM (Fig. 3A–D). MCAM staining indicated continuous populations of MCAM^high^ (spermatogonia) and MCAM^negative^ (spermatocytes) cells, with the latter consisting of three cell populations with different H4V expression levels (P4, P5, and P6; Fig. 3E). P4, P5, P6, and P8 MCAM^low^ cells located between the spermatogonia and spermatocytes (Fig. 3D) were subjected to DNA content analysis and cytological examination. DNA content analysis revealed that P4 cells were 1C (round spermatids), P5 cells were 2C (secondary spermatocytes), and P6 and P8 cells were 4C (primary spermatocytes) (Fig. 3F). Cytological examination by immunostaining with SYCP3 (synaptonemal complex formation) and γH2AX (XY body formation) supported the results of the DNA content analyses (Fig. 3G). Quantification results demonstrated that P6 exhibited >97% late pachytene spermatocytes, while P8 contained ∼95% early pachytene and ∼5% leptotene/zygotene spermatocytes (Fig. 3H). P4 mainly comprised (>95%) round spermatids, while P5 were mainly secondary spermatocytes (∼80%) (Fig. 3H).

**Figure 3.**
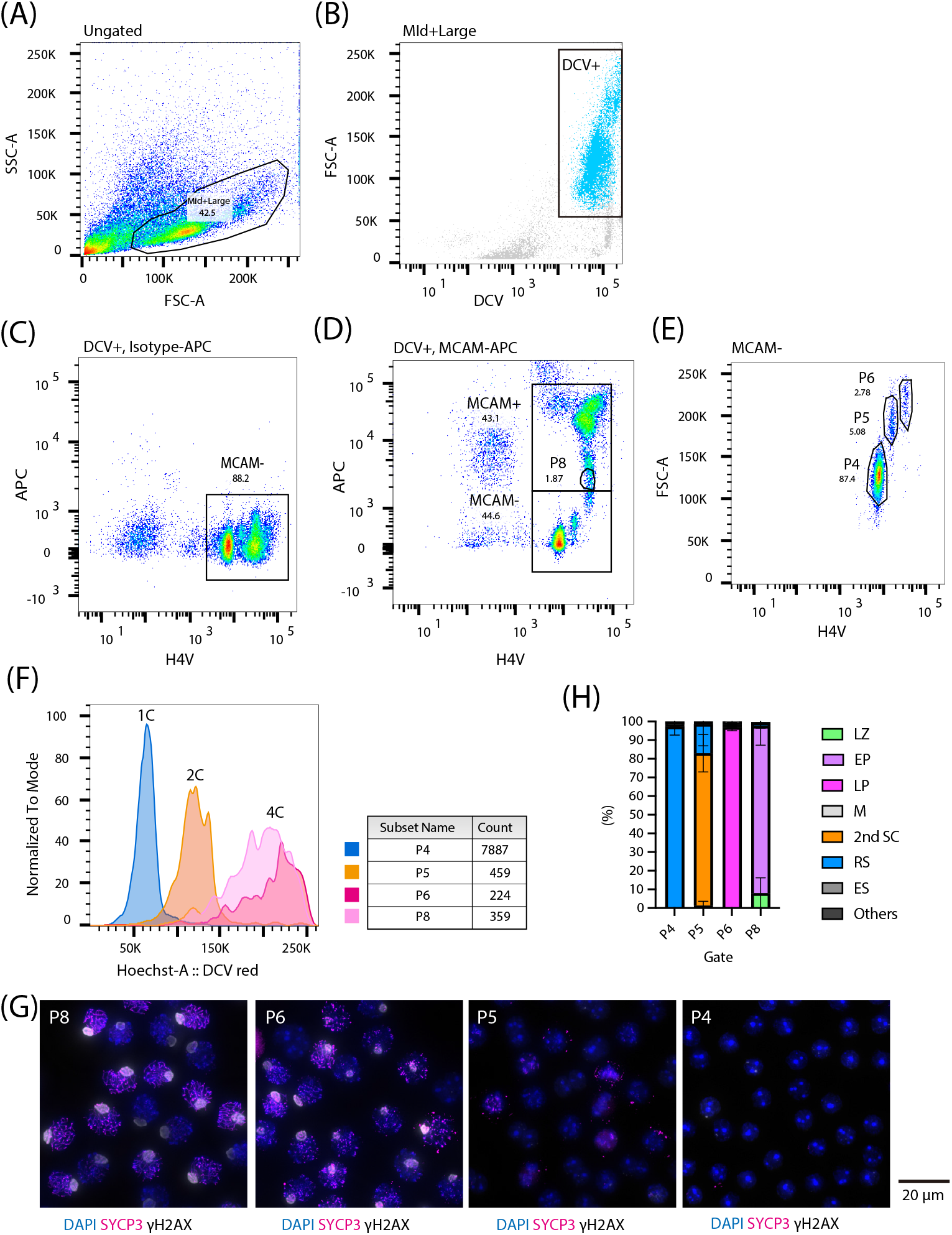
Characterization and purification of meiotic cell subpopulations. (A) Gating strategy to obtain middle and large-sized cells. (B) Gating for DCV+ cells. (C, D) Expression of H4V (X-axis) and MCAM (Y-axis) in DCV+ cells. Staining results with Isotype-APC (C) and MCAM-APC (D) are shown. (E) Gating of three populations (P4, P5, P6) of MCAM-cells. (F) DNA content analyses of the P4–P8 populations. (G) Representative images of immunocytological staining in P4–P8 cell populations. (H) Percentage of meiotic subpopulations and round spermatids in the P4–8 populations.

### Isolating subpopulations of spermatids

Finally, we aimed to separate the post-meiotic spermiogenic cells. In VC mice, the transition from H4-Venus to H3.3-mCherry during spermiogenic steps 9–11 corresponds to the timing of histone-protamine exchange (23). By taking advantage of this unique property, spermatids can be separated according to their spermiogenic steps using the fluorescence intensity and color switching of Venus and mCherry, respectively. For this purpose, small to middle-sized cells were obtained in FSC/SSC (Fig. 4A), followed by singlet and DCV+ cell selection (Fig. 4B, 4C). The VC plot indicated the continuously aligned cells from the V+C-to the V-C+ area via the V+C+ area, similar to that in Fig. 1G, except for depletion of a large cell-derived population (Fig. 4D) and the enrichment of V-C+ spermatids (Fig. 4D). The DCV+ cells were then divided into three populations depending on their approximate cell sizes, considering the dynamic nuclear condensation during spermiogenesis: S1 (very small), S2 (small), and Middle (Fig. 4E). Cell populations of each size were plotted in H4V/H3.3C, and five major cell populations, named P1–P5, were sorted and subjected to cytological analysis (Fig. 4F–H). As expected in Fig. 1, P1 exclusively consisted of round spermatids expressing Venus but not mCherry (Fig. 4I, J). Weak H4 acetylation (H4ac) was also observed in P1 (Fig. 4K). P2 consisted of elongated spermatids that had begun to express mCherry (Fig. 4I, J). H4ac was more intense in some cells, whereas others lost acetylation (Fig. 4K). Based on the staining patterns of PNA and H4ac, we concluded that P1 and P2 consisted of step 1–8 (round) and step 11– 12 (elongated) spermatids, respectively. Sorted P2 cells were subjected to ELSI, and they were able to generate pups with a success rate (21.5%) comparable to a previous study (∼17%) (29), demonstrating the utility of this method (Table 1). In addition, although we previously reported via histological examination that H4-Venus and H3.3-mCherry expression are mutually exclusive, flow cytometry indicated the transient appearance of a double-positive population between P1 and P2, which likely corresponded to step 9–10 spermatids (Fig. 4D, F). PNA and DNA staining identified P3 and P4 as step 13–14 (condensing) and step 15–16 (condensed) spermatids, respectively (Fig. 4I, J). Apparently, P3 cells possessed less cytoplasm than P2 cells did (Fig. 4I). At P4, the tails were clearly visible (Fig. 4I). In contrast, the presence of the P5 population, which was derived from the S1 population, was unexpected (Fig. 4H). The VC plot of this population overlapped with that of P1, despite their smaller sizes (Fig. 4F, H). Cytological examination revealed that the cells at P5 were round spermatids with small, distorted nuclei and missing cytoplasm (Fig. 4I), possibly caused by technical artifacts during the preparation of the single-cell suspension. Stepwise changes in H4V and H3.3C levels during spermiogenesis are summarized in Fig. 5A.

**Figure 4.**
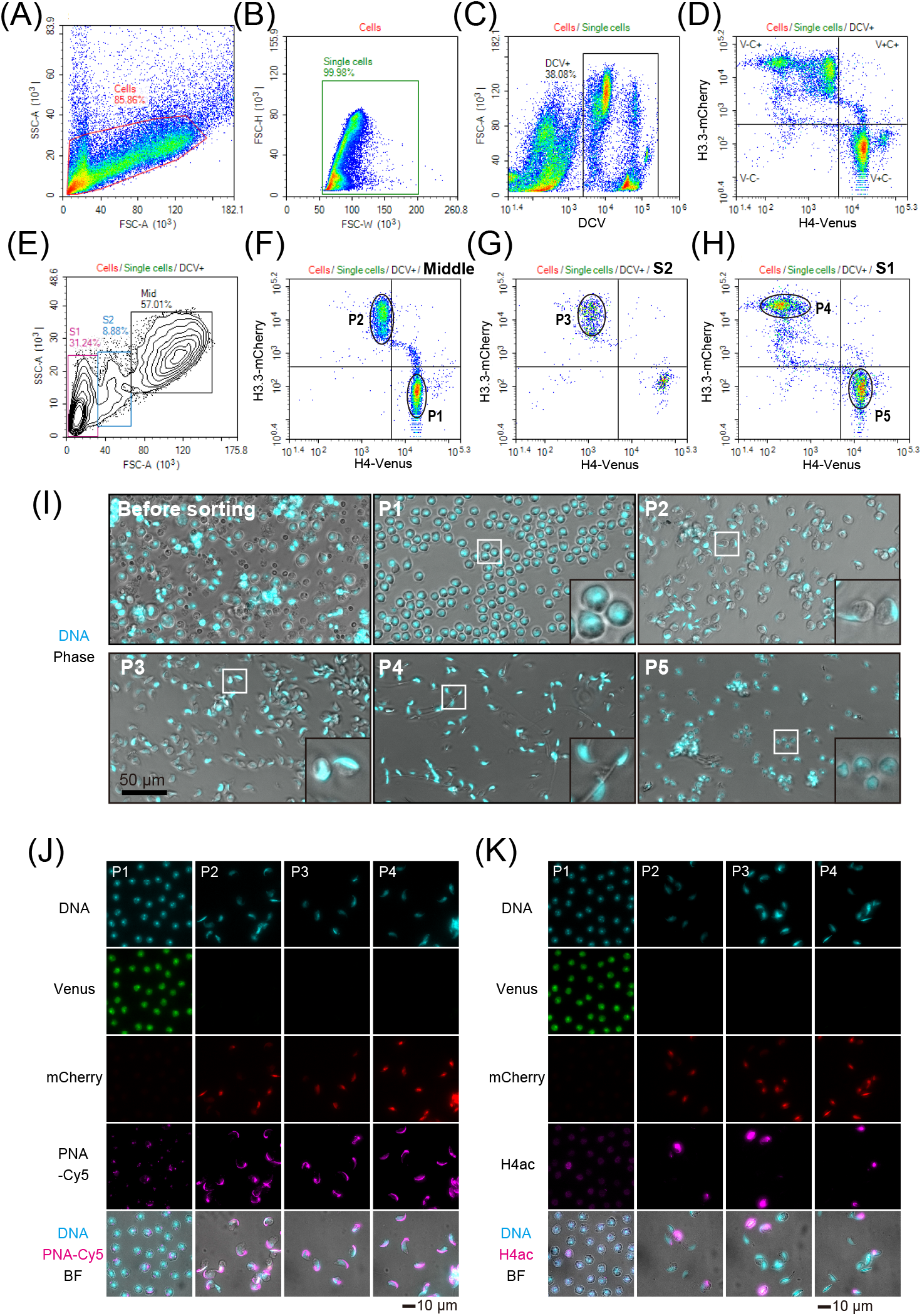
Characterization and purification of spermatid subpopulations. (A) Gating strategy to obtain small and middle-sized cells. (B) Removal of non-single cells. (C) Gating of DCV+ cells. (D) Expression pattern of H4V (X-axis) and H3.3C (Y-axis) in DCV+ cells. (E) Setting of gates according to cell size (S1, S2, and Mid). (F–H) Expression pattern of H4V and H3.3C in middle (F), S2 (G), and S1 (H) cells. P1–P5 gates are indicated. (I) Representative cellular morphologies of P1–P5 cells after cell sorting. (J, K) Representative images of immunocytological staining in P1–P4 populations. BF, bright field.

**Figure 5.**
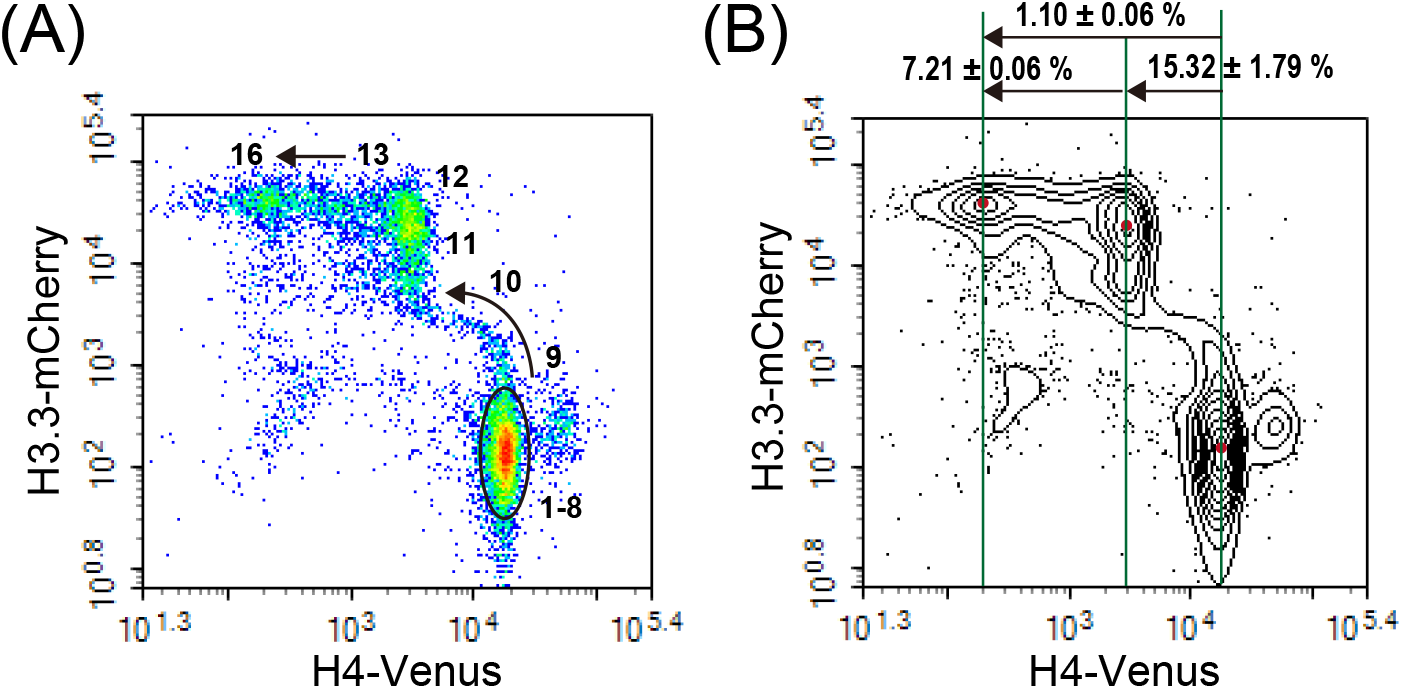
Spermiogenic step-dependent changes in H4V and H3.3C expression. (A) Correspondence between spermiogenic cell differentiation steps and H4V/H3.3C expression. (B) Reduction in H4V levels during spermiogenesis. The degree of reduction in H4V expression is shown as the mean ± SD% (n=3).

**Table 1.** Developmental rate of ELSI-derived embryos

| Number of 1-cell embryos | Number of 2-cell embryos | % of embryos reached to 2-cell stage | No. embryos transferred to surrogate mothers | No. pups born | % of embryos born/transferred |
| --- | --- | --- | --- | --- | --- |
| 428 | 320 | 74.8 | 298 | 64 | 21.5 |

Results from 6 (2-cell rate) and 5 (birth rate) independent experiments are shown.

### Quantification of H4-Venus removal during spermiogenesis

It has been biochemically demonstrated that approximately 1–2% of histone H3 escapes histone-protamine substitution and is retained in the mouse sperm (30). Therefore, we tested whether our flow cytometry data were consistent with these findings. The intensity of H4V reduced to 1.10±0.06% from steps 1–8 to step 16 spermatids, similar to the previous report (Fig. 5B) (30). Furthermore, H4V removal occurred in two steps; first from steps 1–8 to step 12 spermatids (decreased to 15.32±1.79%), and then from step 12 to step 16 (decreased to 7.21±0.06%) (Fig. 5B). It is important to note that the timing of the first removal corresponded to PRM1 incorporation and the second eviction corresponded to PRM2 incorporation (31), which might reflect correlations between histone removal and PRM incorporation.

## Discussion

The purification of testicular germ cells via FACS in combination with appropriate DNA staining dyes is becoming a popular method because of the convenience of obtaining highly purified cells at different spermatogenic stages according to their DNA content. However, the separation of cell subpopulations with the same DNA content, such as meiotic cells (preleptotene, leptotene/zygotene, pachytene, and diplotene) and spermatids (round, elongating, and elongated), is still challenging, as these subpopulations are distributed contiguously on the FACS spectrum. Thus, it is difficult to objectively separate them according to their differentiation steps, even though the differences in their sizes help to some extent. Although the necessity of having transgenes in mice makes them less versatile, the use of VC mice allows the precise separation of these cell subpopulations with high purity. In particular, the transition from Venus to mCherry, which coincides with the histone-protamine exchange, allowed the fine fractionation of spermatids according to their differentiation steps. As various epigenetic events occur stepwise during this period, this method may be ideal for preparing cells to analyze the stepwise changes occurring in these epigenetic events.

This study provides three insights into the chromatin dynamics during spermiogenesis. The first is about histone removal; although H4V is an artificial transgene that accounts for only a small percentage of the total H4 (23), we observed a decrease in H4V levels quantitatively. Interestingly, H4V removal occurs in two distinct steps (spermiogenic steps 8–12 and 12–16), corresponding to when PRM1 and PRM2 are incorporated into the chromatin, respectively (31). Moreover, this may coincide with transient chromatin relaxation in elongating/condensing spermatids (32).

The second is the incorporation of H3.3 at the time of histone removal. A previous study showed that H3.3 is preferentially enriched in elongated spermatids, which may be attributed to newly incorporated H3.3 (30). Although H3.3C is expressed by the Prm1 promoter in the post-meiotic stages, our observations support the idea of a previous study (30). Furthermore, H3.3C was incorporated into spermatids along with a reduction in H4V expression from steps 9–10; the intensity further increased in steps 11–12 without a change in H4V levels (Fig. 5A). Although the Prm1 promoter is possibly more activated in the later steps, this suggests that the incorporation of H3.3 is also regulated in two steps.

The third factor is the fate of the removed histones. We observed a large DCV-negative cell population, presumably consisting of cytoplasmic components called “residual bodies” (Fig. S1). Without DCV staining, this population overlaps with step 11–12 spermatids, accompanied by a decrease in the intensities of both H4V and H3.3C. This observation suggests that some evicted histones translocate to the cytoplasm and are degraded. It has been reported that PA200, a testis-specific proteasome complex, is involved in the degradation of acetylated histones during histone-protamine replacement (33). However, PA200 was shown to function in step 9 spermatids in a histone acetylation-dependent manner, suggesting that the degradation of H4V and H3.3 in residual bodies is PA200-independent. In contrast, the timing of the first decrease in H4V level at steps 9–11 coincides with global histone acetylation and PA200 expression. Although the degradation of H4V and H3.3C may not reflect the behavior of endogenous histones *in vivo*, our results demonstrated that histone degradation happens in two steps, the latter of which likely occurs in the cytoplasm.

Collectively, the use of VC mice for the isolation of testicular germ cells, especially post-meiotic spermatids, enabled fine, step-by-step cell isolation. This method is expected to facilitate various studies on spermatid differentiation, particularly those involving chromatin dynamics.

## Figure Legends

**Figure S1.**
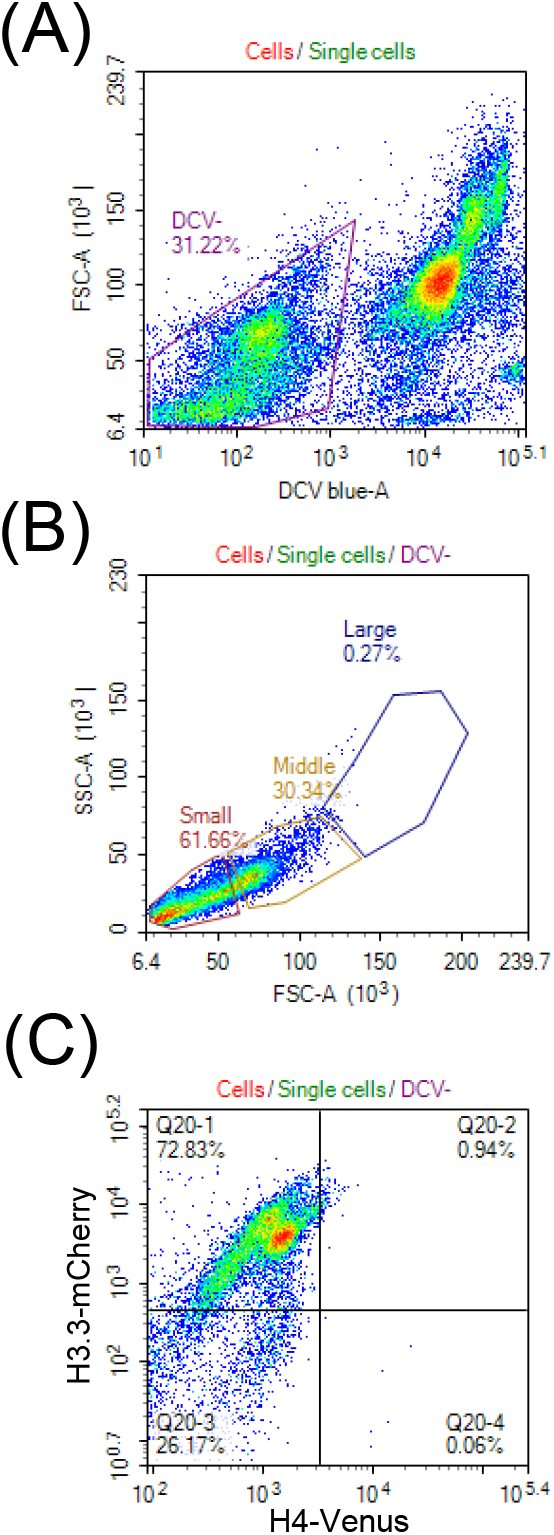
(A) Gating of DCV-negative (DCV-) cells (this plot is the same as Fig. 1C). (B) Size distribution of DCV-cells. The gate setting is the same as in Fig. 1D. (C) Expression pattern of H4V (X-axis) and H3.3C (Y-axis) in DCV-cells.

